# Analysis of receptors responsible for the dysfunction of the human immune system by different viral infections

**DOI:** 10.1101/2022.01.26.477819

**Authors:** Sergey M. Ivanov, Olga A. Tarasova, Vladimir V. Poroikov

## Abstract

There are difficulties in creating direct anti-viral drugs for all viruses, including new, suddenly arising infections, such as COVID-19. Therefore, pathogenetic therapy is often used to treat severe viral infections. Despite significant distinctions in the etiopathogenesis of viral diseases, they are often associated with the substantial dysfunction of the immune system. To identify shared mechanisms of immune dysfunction during infection by nine different viruses (cytomegalovirus, Ebstein-Barr virus, human T-cell leukemia virus type 1, Hepatitis B and C viruses, human immunodeficiency virus, Dengue virus, SARS-CoV, and SARS-CoV-2), we applied analysis of corresponding transcription profiles from peripheral blood mononuclear cells (PBMC). As a result, we revealed common pathways, cellular processes, and master regulators for studied viral infections. We found that all nine viral infections cause immune activation, exhaustion, cell proliferation disruption, and increased susceptibility to apoptosis. An application of network analysis allowed us to identify receptors of PBMC that are the proteins at the top of signaling pathways, which may be responsible for the observed transcription changes. The identified relationships between some of them and virus-induced immune disfunction are new, with little or no information in the literature, e.g., receptors for autocrine motility factor, insulin, prolactin, angiotensin II, and immunoglobulin epsilon.

## Introduction

Viruses cannot replicate without host cells; therefore, their interaction with hosts is crucial for viral existence, replication, and transmission. As opposite, the immune system defends the host organism from the virus. Thus, the development of viral infection and the severity of the disease is mediated by a complex interplay between the virus and host immune system. The intensity and efficacy of the immune response depends on the immune status of a person and the ability of some viruses to disrupt human immunity. Therefore, the peculiarities of interactions between a particular virus and the human body can be a part of a whole pathogenesis process that determines the severity of viral diseases.

Several viruses, e.g., cytomegalovirus, Epstein-Barr virus, human T-cell leukemia virus type 1, hepatitis B and C viruses, human immunodeficiency virus 1, can cause chronic or latent infections. Herpesviruses, including cytomegalovirus and Epstein-Barr virus, cause latent infection and can persist for ages in healthy people with no clinical symptoms, but they can be reactivated by inflammatory stimuli or immunosuppression and cause various pathologies [1,2]. Retroviruses such as human T-cell leukemia virus type 1 and human immunodeficiency virus 1 cause chronic infections by integrating viral genome into human DNA. Human T-cell leukemia virus type 1 usually persists in a human organism without symptoms; however, it may cause adult T cell leukemia/lymphoma or HTLV-1-associated myelopathy/tropical spastic paraparesis in a small part of infected people [3,4]. Human immunodeficiency virus 1 may also persist in the human organism without symptoms for many years but eventually causes acquired immunodeficiency syndrome in almost all infected people [5].

Other viruses from various families, such as coronaviruses, influenza, and Dengue virus, cause acute infection diseases with different degrees of severity. In this case, the disease may lead to death or recovery with the complete elimination of the virus [6,7].

Different viruses have different transmission routes, and target cells and are causes of infectious diseases of different duration and severity. In addition, new, previously unknown viral infections such as COVID-19 may occur; therefore, it is challenging to create direct anti-viral therapeutics due to differences in the structures and functions of viral targets and the long duration of the drug development process. Thus, pathogenetic therapy is often used to treat severe viral diseases. Despite significant distinctions in the etiopathogenesis of viral diseases, many of them cause or increase the dysfunction of the immune system [8–18]. Long-lasting immune response with release of cytokines in the case of chronic infections or cytokine storm in the case of some acute infections, e.g., dengue fever and COVID-19, lead to cytokine-dependent immune activation, which, in turn, causes immune dysfunction or “exhaustion,” significant cell loss and immunodeficiency. As a result, the immune system cannot eliminate viruses effectively, which influences disease severity, duration, and development of secondary infections. For instance, chronic cytokine-dependent activation of T cells by human immunodeficiency virus infection causes their apoptosis with the development of immunodeficiency [19]. Cytomegalovirus increases disease progression and mortality in infants with human immunodeficiency virus infection since it raises activation and apoptosis of both CD4(+) and CD8(+) T cells [11]. COVID-19 and dengue hemorrhagic fever are associated with cytokine storm, which causes the induction of T lymphocytes’ apoptosis, increasing the disease severity [9,17,20]. Thus, it is crucial to identify common pathogenetic mechanisms for different viral infections because it will allow the development of new anti-viral drugs applicable against various infections.

One of the efficient ways to identify common mechanisms of immune dysfunction is a genome-wide analysis of OMICs data [5]. Most of the studies that analyzed virus-host interaction used transcriptomics data obtained by microarray and RNA sequencing approaches [21–29]. Some of these studies focused on comparing transcriptional changes in blood cells caused by the same or similar viral infections, e.g., respiratory viral infections. For instance, Vavougios G.D. analyzed transcriptomics data and determined overlapping host gene signatures between SARS-CoV-2 and other viral and bacterial potential co-pathogens [24]. Dunmire S.K. with colleagues compared the Epstein-Barr transcription response profile to multiple other acute viral infections, including influenza A, respiratory syncytial virus, human rhinovirus, attenuated yellow fever virus, and Dengue virus. They revealed similarity only to Dengue virus profile as well as that of presented in patients with hemophagocytic syndromes, suggesting Epstein-Barr and Dengue viruses cause uncontrolled inflammatory responses [21]. McClain M.T. with colleagues measured transcription profiles from COVID-19 patients and directly compared them to subjects with seasonal coronavirus, influenza, bacterial pneumonia, and healthy controls. They identified a 23-gene signature, which can be utilized to diagnose and differentiate COVID-19 from other viral diseases [27].

The current study aimed to analyze the transcriptomics data for viral diseases causing or increasing immune dysfunction. We compared the transcription profiles in blood cells between pathologies caused by nine viruses. They all cause immune dysfunction and immunodeficiency, and the corresponding transcriptomics data, measured at the same conditions, including cell types, are available in public databases. We identified common differentially expressed genes (DEGs), pathways, and master regulators, which are the proteins at the top of signaling pathways responsible for the observed transcription changes. Particularly, we found receptors for cytokines, growth factors, hormones, and mediators responsible for immune dysfunction in most studied viral diseases.

## Materials and Methods

### Collection of transcription datasets

We performed a comprehensive search across two transcriptomics databases: Gene Expression Omnibus (GEO) [30], and ArrayExpress [31] using the following query: “(virus OR viral) AND (lymphocytes OR “B cells” OR “T cells” OR monocytes OR “NK cells” OR PBMC OR “peripheral blood mononuclear” OR neutrophils OR “whole blood”).” We selected datasets containing data on gene transcription in blood cells measured by microarrays or RNA sequencing and obtained from both viral-infected and uninfected people. We selected only datasets obtained from people who did not take anti-viral therapeutics. In total, we identified 112 data sets, which were different by the virus, cell type, and experimental method. The most frequently used cell type was peripheral blood mononuclear cells (PBMC). The datasets related to these cells contain transcription profiles from patients infected by SARS-CoV-1, SARS-CoV-2, Dengue virus, human immunodeficiency virus 1, human T lymphotropic virus type 1, hepatitis B and C viruses, cytomegalovirus, and Epstein–Barr virus.

### Genome Enhancer pipeline

To identify differentially regulated genes and master regulators (MRs), we used the Genome Enhancer tool [32] developed by geneXplain GmbH [33]. Briefly, Genome Enhancer implemented a pipeline including four main steps:

1. Normalization of gene transcription data and identification of differentially expressed genes (DEGs);
2. Identification of the regulatory regions (promoters and enhancers) of the DEGs;
3. Analysis of the regulatory regions of DEGs to predict transcription factor binding sites (TFBSs), identify those enriched in promoters and enhancers of DEGs compared to genes with unchanged transcription, and predict the combinations of TFBSs called “composite regulatory modules,” which may reflect complexes of co-acting TFs;
4. Reconstruction of the signaling pathways that activate these TFs and identify MRs at the top of such pathways.

To identify receptors among revealed MRs, we retrieved a list of receptors from the OmniPath database [34] using the OmnipathR R package.

### Pathway enrichment analysis

To identify KEGG pathways [35] that were differentially regulated in PBMC from a particular viral infection-related group of patients compared to healthy control, we peformed pathway enrichment analysis [36] using the “enrichr” function from the “enrichR” R package [37]. Function “enrichr” allows identifying pathways “enriched” with differentially regulated genes compared to the random gene sets. We performed the corresponding analysis for up- and down-regulated genes separately. We selected pathways associated with at least three up- or down-regulated genes with a p-value of less than 0.05, which are widely accepted thresholds.

### Comparison of KEGG pathways, functionally related groups of genes, and master regulators between viral infection diseases

To compare lists of KEGG pathways, manually selected groups of genes, and MRs obtained for infectious diseases caused by nine viruses, we created heatmaps using the “pheatmap” R package [38].

## Results

### Data on viral-related gene transcription in peripheral blood mononuclear cells

We performed a comprehensive search across transcriptomics databases and selected 11 transcription datasets from the Gene Expression Omnibus (GEO) database containing data on gene transcription in peripheral blood mononuclear cells (PBMC) from healthy people and patients infected by one of the nine viruses (see Table 1). The details are provided in the Materials and Methods section. Some datasets contain samples from patients with a different form of viral infection disease, e.g., different severity (dengue fever or dengue hemorrhagic fever, severe or moderate COVID-19), or duration of disease (acute or latent Epstein-Barr and cytomegalovirus infections). Further analysis was performed separately for each of these viral infection-related groups. We defined “viral infection-related group” as a group of samples obtained from a particular transcription dataset and related to the particular viral infection in a particular form (e.g., severe or moderate, acute or chronic, symptomatic or asymptomatic).

**Table 1.**
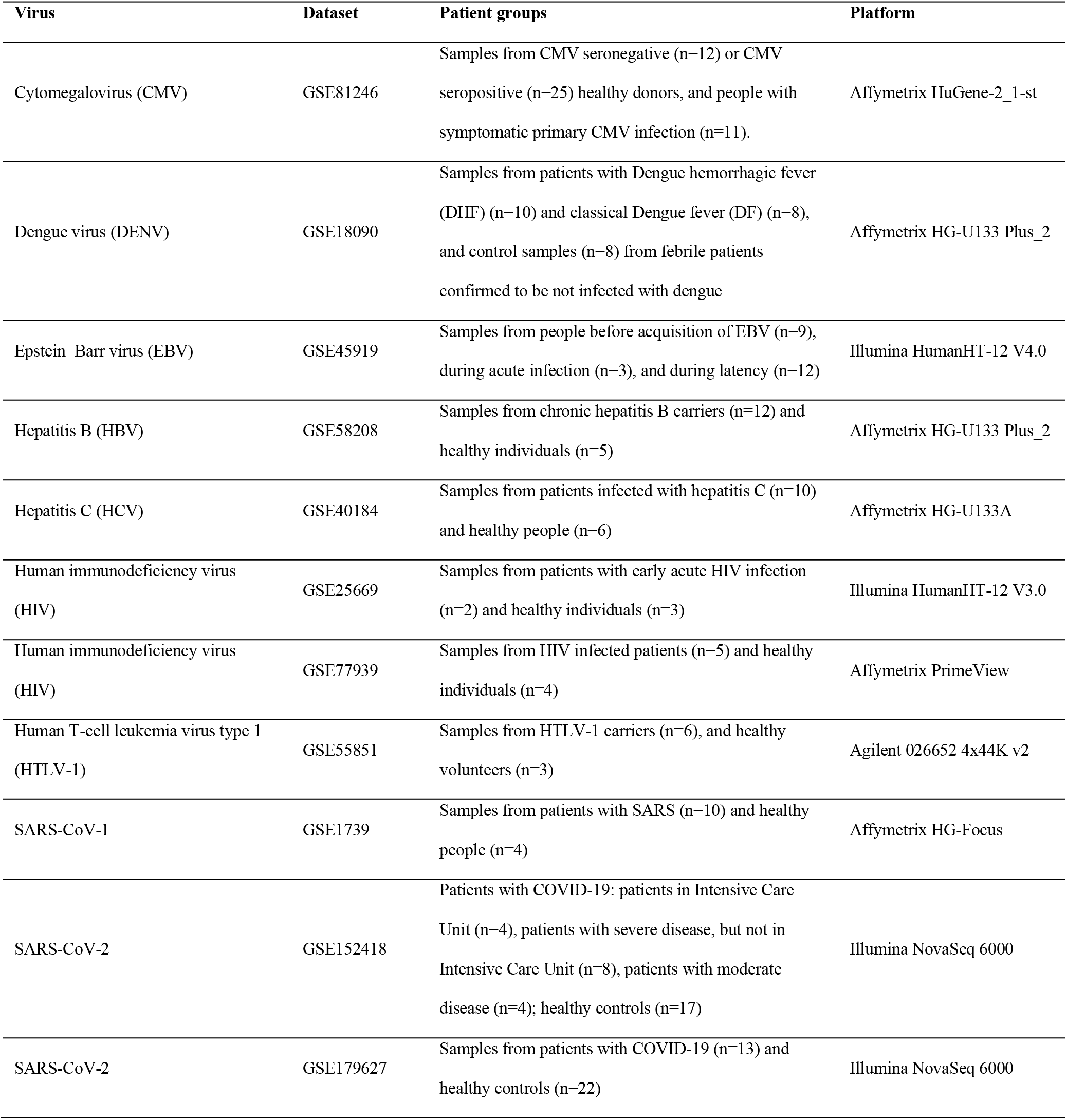
Transcription datasets with information on gene transcription in PBMC derived from healthy people and patients infected by one of 9 viruses.

| Virus | Dataset | Patient groups | Platform |
| --- | --- | --- | --- |
| Cytomegalovirus (CMV) | GSE81246 | Samples from CMV seronegative (n=12) or CMV seropositive (n=25) healthy donors, and people with symptomatic primary CMV infection (n=11). | Affymetrix HuGene-2_1-st |
| Dengue virus (DENV) | GSE18090 | Samples from patients with Dengue hemorrhagic fever (DHF) (n=10) and classical Dengue fever (DF) (n=8), and control samples (n=8) from febrile patients confirmed to be not infected with dengue | Affymetrix HG-U133 Plus_2 |
| Epstein-Barr virus (EBV) | GSE45919 | Samples from people before acquisition of EBV (n=9), during acute infection (n=3), and during latency (n=12) | Illumina HumanHT-12 V4.0 |
| Hepatitis B (HBV) | GSE58208 | Samples from chronic hepatitis B carriers (n=12) and healthy individuals (n=5) | Affymetrix HG-U133 Plus_2 |
| Hepatitis C (HCV) | GSE40184 | Samples from patients infected with hepatitis C (n=10) and healthy people (n=6) | Affymetrix HG-U133A |
| Human immunodeficiency virus (HIV) | GSE25669 | Samples from patients with early acute HIV infection (n=2) and healthy individuals (n=3) | Illumina HumanHT-12 V3.0 |
| Human immunodeficiency virus (HIV) | GSE77939 | Samples from HIV infected patients (n=5) and healthy individuals (n=4) | Affymetrix PrimeView |
| Human T-cell leukemia virus type 1 (HTLV-1) | GSE55851 | Samples from HTLV-1 carriers (n=6), and healthy volunteers (n=3) | Agilent 026652 4x44K v2 |
| SARS-CoV-1 | GSE1739 | Samples from patients with SARS (n=10) and healthy people (n=4) | Affymetrix HG-Focus |
| SARS-CoV-2 | GSE152418 | Patients with COVID-19: patients in Intensive Care Unit (n=4), patients with severe disease, but not in Intensive Care Unit (n=8), patients with moderate disease (n=4); healthy controls (n=17) | Illumina NovaSeq 6000 |
| SARS-CoV-2 | GSE179627 | Samples from patients with COVID-19 (n=13) and healthy controls (n=22) | Illumina NovaSeq 6000 |

We identified up- and down-regulated genes for each viral infection-related group (Table 2) with absolute values of log fold changes exceeded 0.5 and p-values less than 0.05. These thresholds were selected empirically to balance the numbers of DEGs and their statistical significance.

**Table 2.** Numbers of up- and down-regulated genes associated with viral infection-related groups.

| Sample group | Abbreviation | Up-regulated | Down-regulated |
| --- | --- | --- | --- |
| Cytomegalovirus primary infection | CMV_pr | 833 | 943 |
| Cytomegalovirus seropositive healthy donors | CMV_pos | 8 | 8 |
| Ebstein-Barr acute infection | EBV_a | 2343 | 2373 |
| Ebstein-Barr latent infection | EBV_l | 3255 | 3315 |
| Hepatitis B chronic infection | HBV | 936 | 1097 |
| Hepatitis C chronic infection | HCV | 136 | 308 |
| Human T-cell leukemia virus type 1 infection | HTLV_1 | 430 | 495 |
| Human immunodeficiency virus infection (GEO ID: GSE25669) | HIV_1 | 333 | 457 |
| Human immunodeficiency virus infection (GEO ID: GSE77939) | HIV_2 | 192 | 108 |
| Dengue hemorrhagic fever | DHF | 878 | 608 |
| Dengue fever | DF | 384 | 368 |
| Severe acute respiratory syndrome | SARS | 720 | 836 |
| Coronavirus Disease 2019 (COVID-19) - patients with evident clinical symptoms | COVID19 | 1605 | 2004 |
| Coronavirus Disease 2019 (COVID-19) - patients in Intensive Care Unit (ICU) | COVID19_ICU | 2925 | 2640 |
| Coronavirus Disease 2019 (COVID-19) - patients with severe disease, but not in Intensive Care Unit (ICU) | COVID19_Sev | 2104 | 1698 |
| Coronavirus Disease 2019 (COVID-19) - patients with moderate disease | COVID19_Mod | 1699 | 1562 |

We considered a gene up- or down-regulated by a particular viral infection if it was up- or down-regulated in at least one of viral infection-related groups, e.g., dengue fever or dengue hemorrhagic fever. We found no genes that were up- or down-regulated by all nine or eight viral infections. There were 1, 8, 31, and 104 up-regulated genes, and 0, 5, 27, and 134 down-regulated genes associated with exactly seven, six, five and four viral infections. The observed small intersections between DEGs may be explained by differences in the pathogenesis of viral infections and differences in experimental methods, microarray platforms, and batch effects. We suggested analyzing the similarity of viral infections at the level of biological pathways.

### Identification for biological pathways that are common for different viral infections

We performed a pathway enrichment analysis (see Materials and Methods) to identify KEGG pathways [35], which are up- or down-regulated in PBMC during studied viral infections. We considered a pathway differentially regulated by a particular viral infection if it was differentially regulated in at least one viral infection-related group regardless of the direction of regulation (up- or down-regulation). We selected pathways, which were differentially regulated by at least half of viral infections, for further analysis (Fig 1).

**Fig 1.**
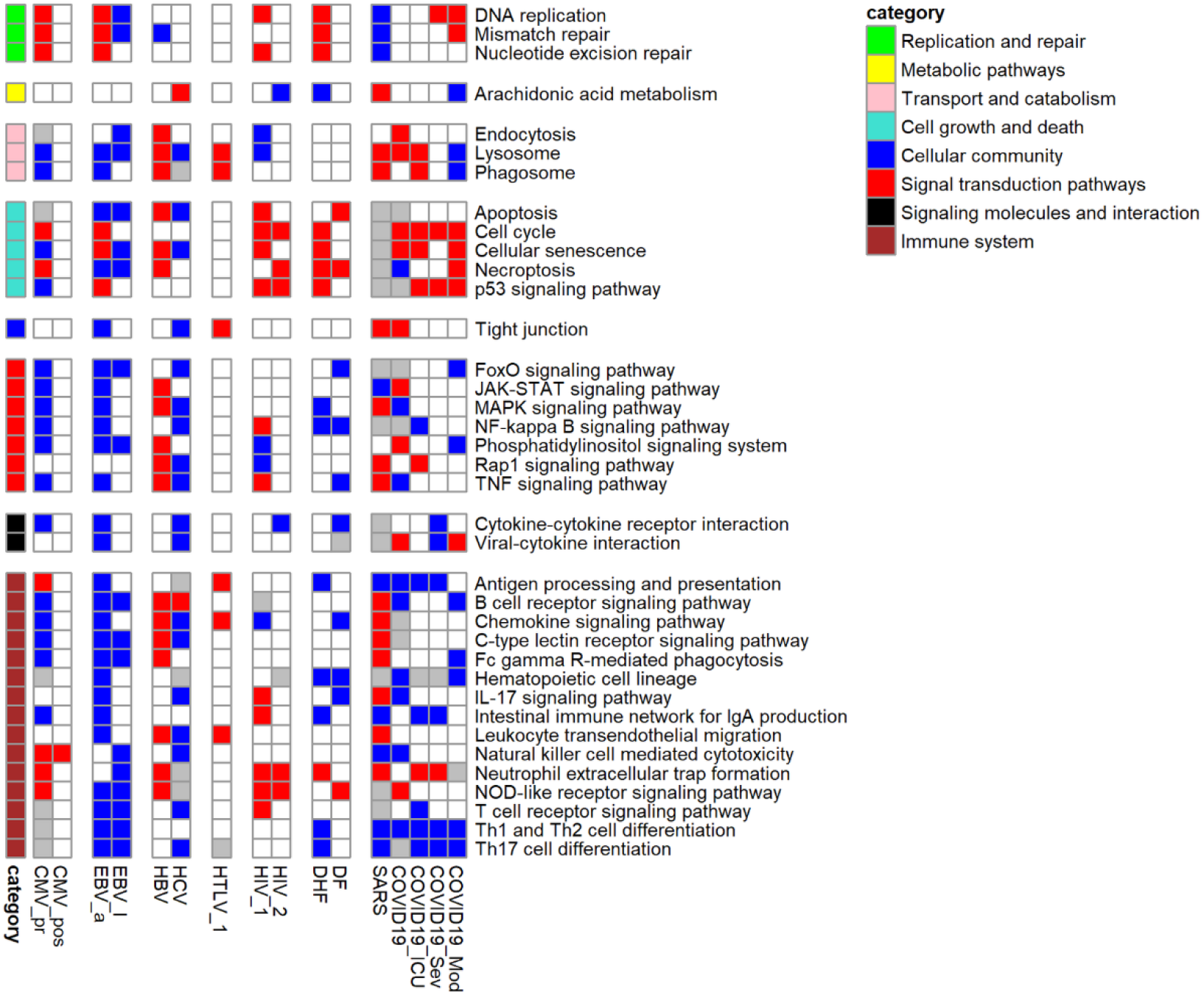
KEGG pathways, up- or down-regulated in PBMC by at least five viral infections. The red color of the cell means that the pathway is enriched by up-regulated genes, the blue color of the cell means that the pathway is enriched by down-regulated genes, grey color of the cell means that the pathway is enriched by both up- and down-regulated genes.

We manually investigated the positions of DEGs in the pathway maps and selected groups of functionally similar genes, which characterize the functional states of PBMC in all nine viral infections (Fig 2).

**Fig 2.**
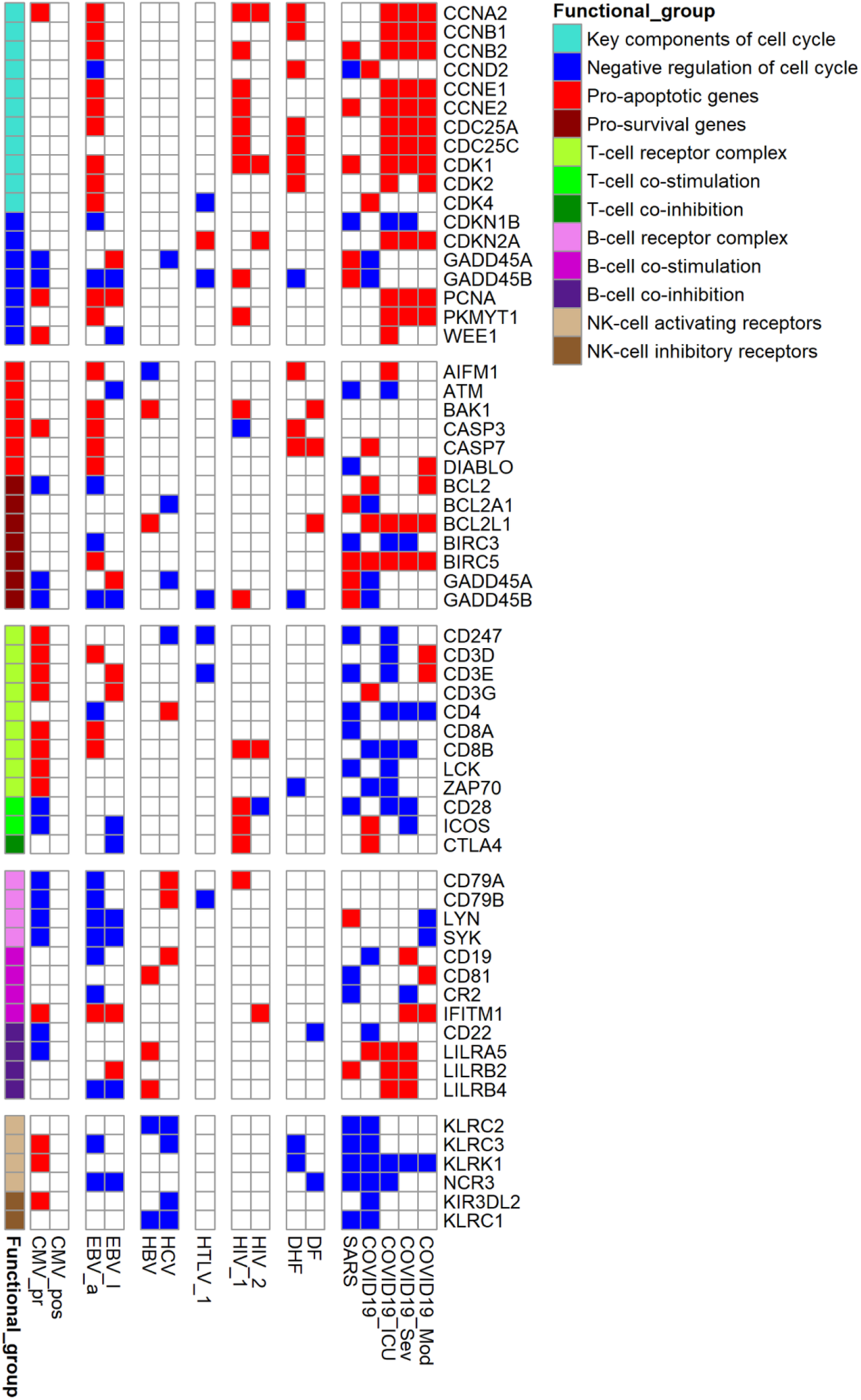
Functionally related groups of genes, up- or down-regulated in PBMC by at least three viral infections. The red color of the cell means that the gene is up-regulated, the blue color of the cell means that the gene is down-regulated. The process is a cell process related to the functional state of PBMC. Regulation is a positive or negative role of the gene in controlling function of the corresponding process.

Figs 1 and 2 demonstrate that PBMC of patients with the most studied viral infections are associated with impaired proliferation, specific immune functions, and increased apoptosis.

### Identification for receptors - master regulators that are common for different viral infections

We identified master regulators (MRs) using the Genome Enhancer pipeline (see Materials and Methods) (Fig 3).

**Fig 3.**
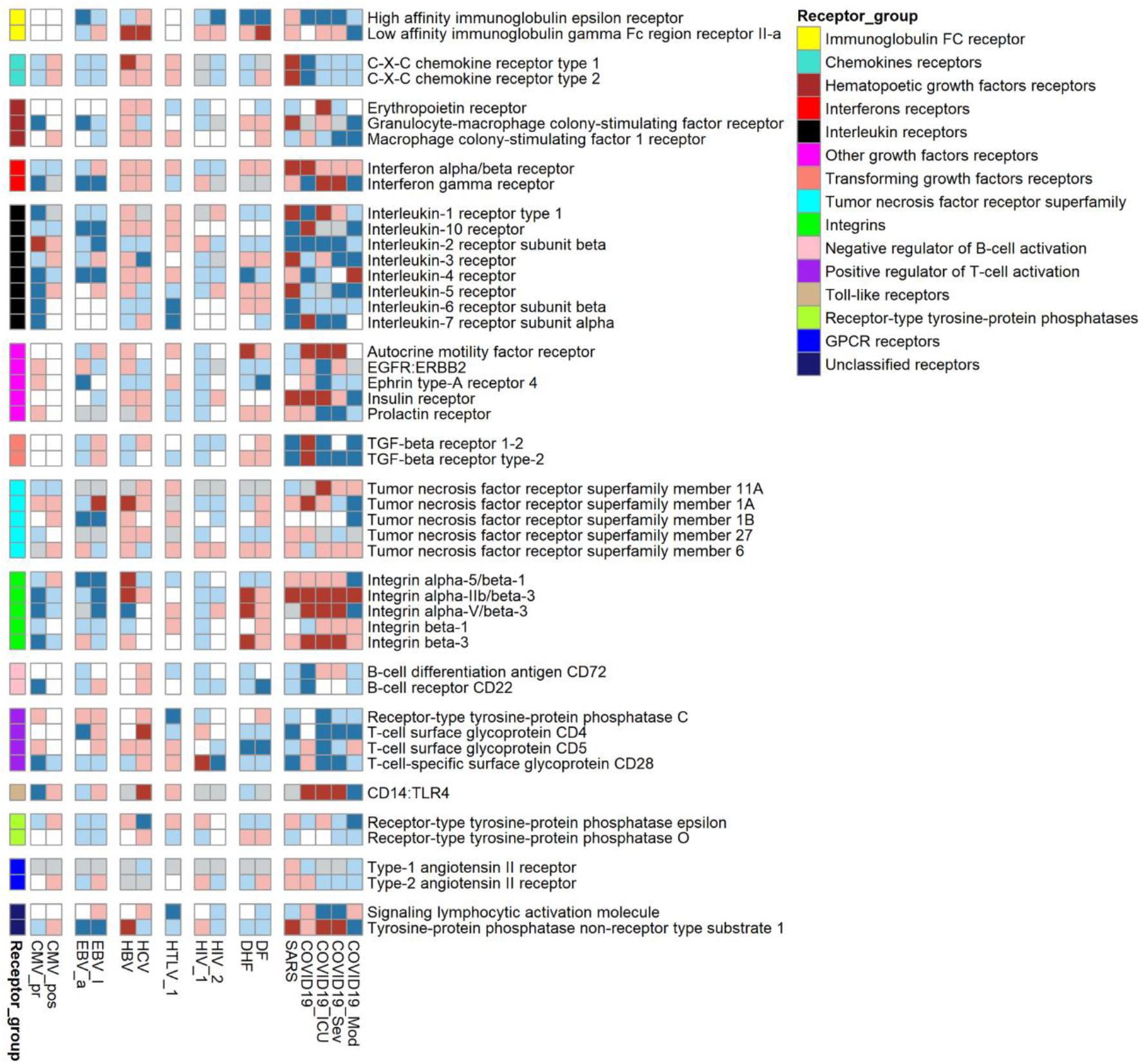
Receptors were identified as MRs for observed transcription changes in PBMC for at least seven viral infections. The red color of cells means that the MR is responsible for the observed up-regulation of genes, dark red color means that the MR is responsible for the observed up-regulation of genes and is up-regulated itself, blue color means that the MR controls the observed down-regulation of genes, dark blue color means that the MR is responsible for the observed down-regulation of genes and down-regulated itself, grey color means that the MR controls both up- and down-regulation of genes.

MRs are the proteins at the top of the signaling network regulating the activity of transcription factors and their complexes and, in turn, are responsible for expression changes observed in PBMC. Most obtained MRs are intracellular “hubs,” such as kinases, phosphatases, ubiquitin ligases, GTPases, and transcription factors. We focused on receptors because their interaction with corresponding ligands is the first of the consequent events leading to gene transcription changes. We selected top receptors identified as MRs for at least seven of nine viral infections (Fig 3). As a result, correspondingly, 18, 9, and 20 receptors were common for seven, eight and nine viral infections. The receptors were grouped according to their belonging to families and functional similarities. Fig 3 also demonstrates that MRs are up- or down-regulated themselves in part of viral infection-related groups.

## Discussion

To identify common molecular mechanisms associated with immune dysfunction caused by both similar and different viral infections, we compared the transcription profiles in PBMC between pathologies caused by nine viruses: SARS-CoV-1, SARS-CoV-2, Dengue virus, human immunodeficiency virus 1, human T lymphotropic virus type 1, hepatitis B and C viruses, cytomegalovirus, and Epstein–Barr virus. These viruses are known to cause immune dysfunction and immunodeficiency. We estimated corresponding mechanisms at the level of biological pathways (Fig 1), smaller groups of function-ally related genes (Fig 2), and receptors, the master regulators that are the proteins at the top of signaling pathways responsible for the observed transcription changes.

First, we identified DEGs for each viral infection-related group reflecting disease severity (dengue fever or dengue hemorrhagic fever, severe or moderate COVID-19) or duration (acute or latent Epstein-Barr and cytomegalovirus infections). There were hundreds or thousands DEGs for all groups except latent cytomegalovirus infection that may indicate significant immune dysfunction in the corresponding pathologies.

Pathway enrichment analysis allowed us to identify KEGG pathways associated with each viral infection-related group (Fig 1). Through the manual analysis of positions of DEGs in the corresponding pathway maps, we identified the small groups of functionally related genes responsible for the regulation of the cell cycle, apoptosis, and immune cell activation and exhaustion (Fig 2). Figs 1 and 2 demonstrate that all viral infections are associated with changes in these processes. The obtained results follow known data on the immunopathogenesis of studied viral infections. For instance, primary cytomegalovirus infection is associated with transcriptional changes that reflect the functional states of immune cells. The transcription of the caspase 3 gene is increased, whereas the transcription of pro-survival genes BCL2, GADD45A, and GADD45B are decreased, which may indicate the PBMC have a pro-apoptotic phenotype. The genes encoding subunits of the T-cell receptor complex are upregulated, but genes encoding co-stimulation receptors CD28 and ICOS are down-regulated, which may indicate the presence of immune activation with disruption of T-cell functions. Primary cytomegalovirus infection is known to cause lymphocyte activation and apoptosis, lymphocytosis, and immune exhaustion [10,11,39]. Human immunodeficiency virus infection is associated with the up-regulation of cyclins A, B, and E. Since PBMC are asynchronized in cell cycle phases, this observation may indicate cell cycle arrest in the G2 phase [40]. It is known that viral protein vpr is associated with G2 arrest in both infected and uninfected blood cells [41]. Down-regulation of co-stimulation receptor CD28 and increased transcription of immune checkpoint CTLA4 indicates that human immunodeficiency virus infection causes activation and exhaustion of T-lymphocytes. It is well known that the long-lasting activation of T-cells by chronic infections, including human immunodeficiency virus infection, leads to the development of their “exhausted” state, which is characterized by increased expression of immune checkpoints, disrupted proliferation, decreased functions, and increased apoptosis [14,40]. Hepatitis C can disrupt the functions of NK cells, which is associated with chronic hepatitis C [42]. The chronic inflammation and immune activation leading to the exhaustion of immune cells is also related to hepatitis C infection [43].

Since the observed transcriptional changes are associated with well-known immunopathogenesis of studied viral infections, we searched for master-regulators, which are the proteins at the top of signaling networks and are responsible for the observed transcriptional changes. Our study focused on receptors because their interaction with corresponding ligands is the first of the consequent events leading to gene transcription changes. We identified many cytokines and growth factors receptors, which are common for studied viral infections (Fig 3). According to Human Proteins Atlas data [44], all of them are expressed in PBMC: lymphocytes, monocytes, or NK cells. The revealed receptors are up- or down-regulated themselves in part of viral infection-related groups (Fig 3). It possibly means that they are part of positive feedback loops and are extremely important to maintain the observed transcription profiles.

The revealed receptors differ in the novelty of their relationships with the functional states of studied immune cells. For instance, the receptors for interferons alpha and gamma, and interleukins 1-10, have a clear, well-known relationship with functional states of PBMC. They are essential for cell activation, differentiation, survival, and proliferation. The receptors like immunoglobulin gamma and epsilon receptors and C-X-C chemokine receptors type 1 and 2 (receptors for interleukin 8) are commonly associated either with other immune cell types functions, or their functions in immune cells are poorly described. For example, CXCR1 and CXCR2 genes encode receptors for interleukin 8, which is well known as a potent neutrophil chemotactic factor. However, the CXCR1 receptor is also expressed on the cell surface of T cells, and interleukin 8 treatment results in a reduction in the activation status of both effector memory T cells and terminally differentiated effector T cells [45]. Interleukin 8 is also an essential chemotactic factor for NK cells [46]. Reduced expression of CXCR1 is associated with decreased counts of NK cells in lymph nodes and incomplete control of HIV viral replication during the early stages of the disease [47]. The FCGR2A gene encodes low-affinity immunoglobulin gamma Fc region receptor II-a that binds to the Fc region of immunoglobulins gamma and promotes phagocytosis of opsonized viral particles [48,49]. Polymorphisms in the FCGR2A gene are associated with severity and outcome of viral infections such as COVID-19 [50], Influenza A [51], and HIV infection [52,53]. Surprisingly, we also identified high-affinity immunoglobulin epsilon receptor, which is well known to be expressed in mast cells and basophils and is responsible for initiating the allergic response. However, it was recently shown that CD4(+) T cells also express immunoglobulin epsilon receptors. Immunoglobulin epsilon induces CD4(+) T-cell production of interleukin 6 and interferon-gamma but reduces their production of interleukin 10 [54]. Polymorphism in the FCER1A gene, which encodes the alpha subunit of the receptor, is associated with the efficacy of hepatitis B treatment by peginterferon alfa-2a [55].

Some of the revealed receptors are not specific to the immune system, and their associations with immune response are usually not well studied. For instance, the role of Ephrin type-A receptor 4 is well described for the nervous system; however, it is also expressed on memory T cells and is responsible for their migration [56]. The insulin receptor is essential for normal adaptive immune function through modulating T cell nutrient uptake and associated glycolytic and respiratory capacities [57]. Moreover, following insulin binding, the insulin receptor translocates to the nucleus, where it plays a crucial role in regulating the transcription of various immune-related genes, including pathways involved in viral infections. Therefore, diabetes with insulin resistance can directly contribute to an inadequate immune response, e.g., observed during COVID-19 [58]. Prolactin and its receptors play an important role in regulating innate and adaptive immune responses [59], particularly in T lymphocyte growth and activation [60,61]. Stimulation of angiotensin AT1 receptors is important for T cell activation and adhesion/transmigration through the basal endothelial membrane, whereas angiotensin AT2 receptors limit this effect [62,63]. AMFR gene encodes the receptor for autocrine motility factor, the protein with multiple functions. It is little known about its functions in the immune cell. Particularly, the autocrine motility factor stimulates immunoglobulin secretion by cultured human PBMC [64].

The role of the most receptors mentioned above in infectious diseases is described for few or none of the studied viruses. Thus, identified common associations between viral infections and the receptors, especially ephrin, prolactin, angiotensin, and autocrine motility factor receptors, are novel and may be a basis for further research. Particularly, the revealed receptors may be investigated as potential targets for treating severe viral infections.

## Conclusions

Since many different viruses currently exist and new viral infections, such as COVID-19, suddenly become spreading among human population, it is challenging to create direct therapeutics for each virus; therefore, pathogenetic therapy is often used to treat severe viral infections. Even though significant differences between viral diseases exist, they are usually associated with the dysfunction of the immune system and immunodeficiency. As a result, the immune system cannot effectively eliminate viruses, which increase disease severity and duration and susceptibility to secondary infections. Thus, it is crucial to identify immunopathogenic mechanisms common for different viral infections. In our study, we applied network-based transcriptomics analysis to identify mechanisms of immune dysfunction caused by nine different viral infections at the levels of pathways, cellular processes, and master regulators, which are the key proteins responsible for the observed immune states. Analysis of revealed pathways and small groups of functionally related differentially expressed genes demonstrated that nine studied viral infections cause immune activation, exhaustion, cell proliferation disruption, and increased susceptibility to apoptosis for peripheral blood mononuclear cells. Since the pathways and processes revealed from the transcriptional changes are associated with known data from the literature, we performed a search for receptors – master regulators, which are at the top of immune cells’ signaling networks and may be responsible for the observed transcriptional changes, and, thus, accountable for the observed immune dysfunction. Besides well-characterized receptors for interleukins and interferons, we identified receptors whose relationships with virus-induced immune disfunction are new, with little or no information in the literature. The receptors for autocrine motility factor, insulin, prolactin, angiotensin II, and immunoglobulin epsilon may be essential for the normal functioning of immune cells and involved in their dysfunction during different viral infections. The revealed receptors may be investigated as potential targets for treating severe viral infections.

## Acknowledgments

We are grateful to the geneXplain GmbH for providing access to the Genome Enhancer pipeline available online at https://ge.genexplain.com.

